## Supplementary Figures and Tables for "Coupling of H3K27me3 recognition with transcriptional repression through the BAH-PHD-CPL2 complex in *Arabidopsis*"

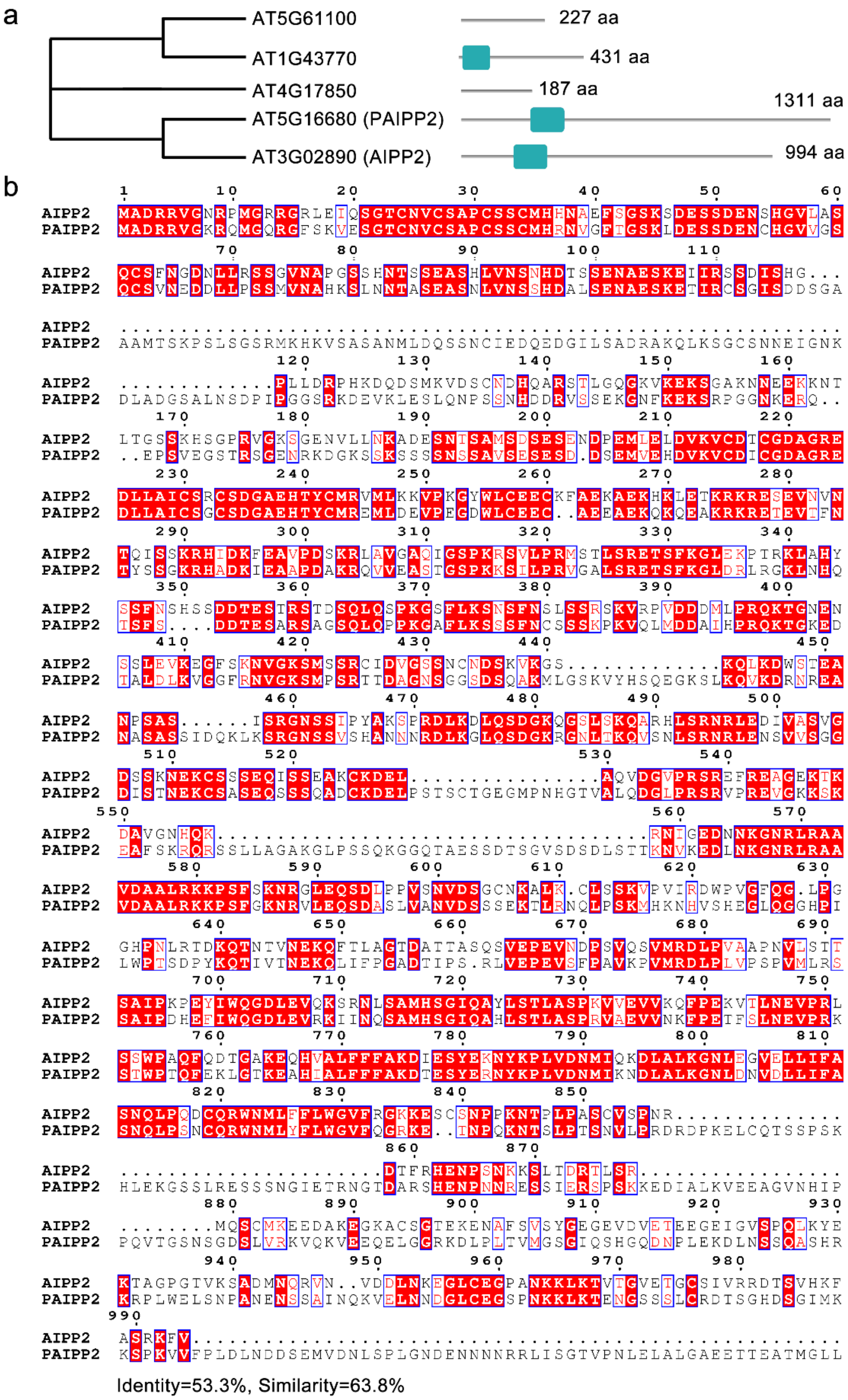


**Supplementary Fig. 1 PAIPP2 is the closest paralog of AIPP2 in *Arabidopsis*.**
**a,** The phylogenetic tree between AIPP2 and its paralog proteins in *Arabidopsis* (left panel) and their domain structures. Green boxes indicate PHD domains. **b,** The sequence alignment between AIPP2 and PAIPP2 showing the high similarity.

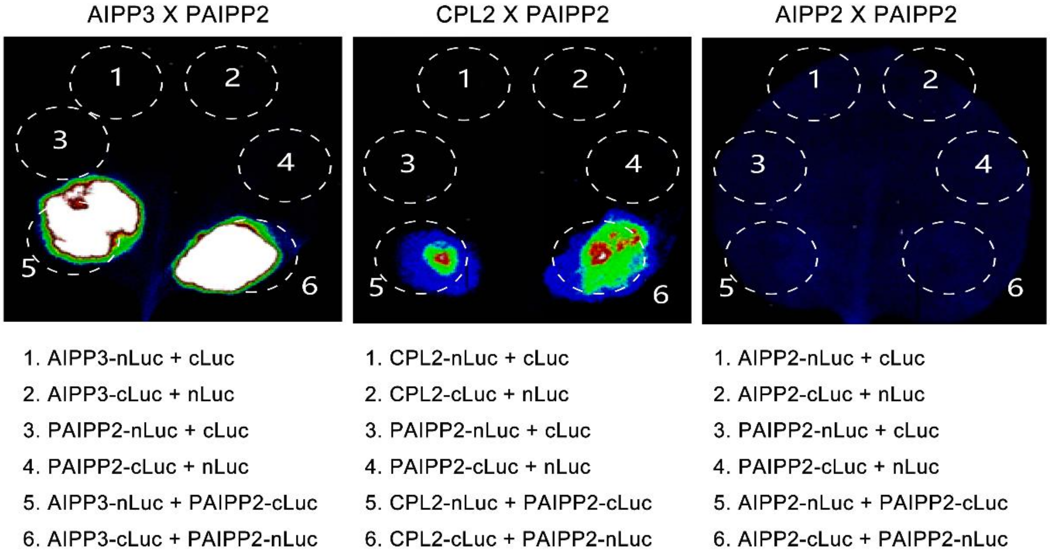


**Supplementary Fig. 2 Protein interactions revealed by split luciferase assay**

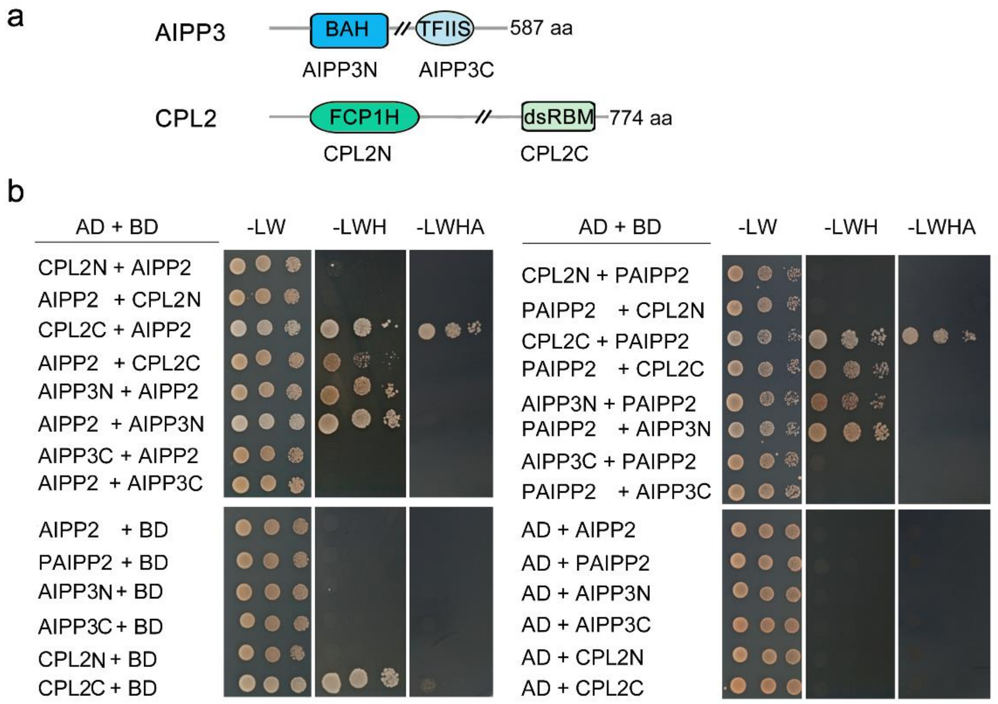


**Supplementary Fig. 3 Domain requirements for the interactions between BAH-PHD-CPL2 complex proteins. a,** The diagram showing the domains and truncated forms of different proteins. **b,** Y2H assays showing the interactions between the truncated forms of proteins. N and C represent the N terminus and C terminus of the proteins, respectively.

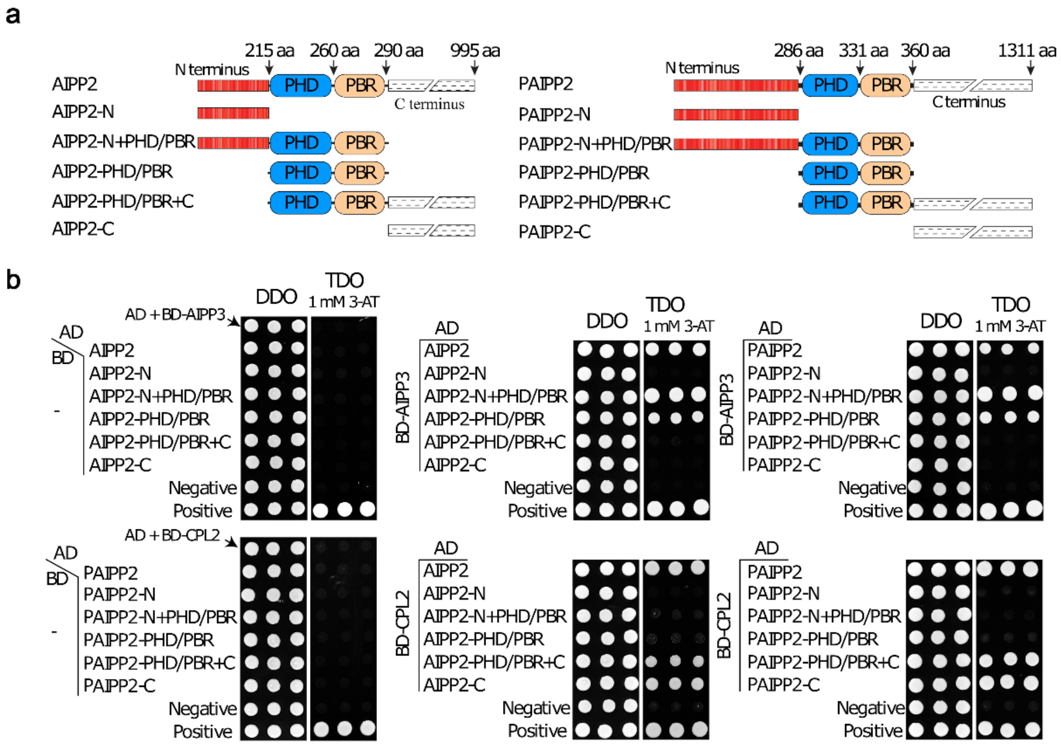


**Supplementary Fig. 4 Domain requirement for protein interactions. a,** Diagrams showing the split domain structure for protein interactions in b and c. **b,** Yeast two-hybrid results showing the protein interactions of different combinations.

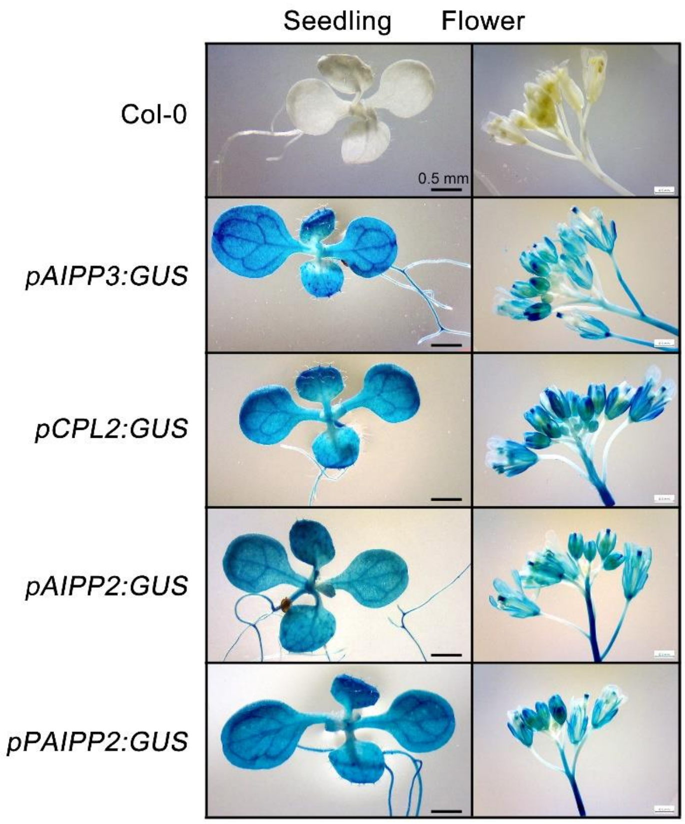


**Supplementary Fig. 5 Expression analysis of the *GUS* reporters from BAH-PHD-CPL2 complex genes in seedlings and inflorescence tissues.** *GUS* reporter genes were expressed in transgenic *Arabidopsis* under the direction of native *AIPP3*, *AIPP2*, *PAIPP2* and *CPL2* promoters. The seedlings and inflorescence tissues were stained to detect the GUS activity.

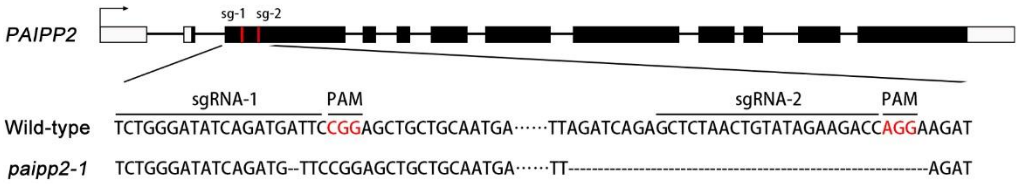


**Supplementary Fig. 6 CRISPR/Cas9-mediated mutagenesis of *PAIPP2*.** The *paipp2-1* was generated by CRISPR/Cas9-mediated mutagenesis. Two sgRNAs were designed to target the N-terminal exon of *PAIPP2*. The *paipp2-1* mutation contains one nucleotide deletion and 33 bp deletion.

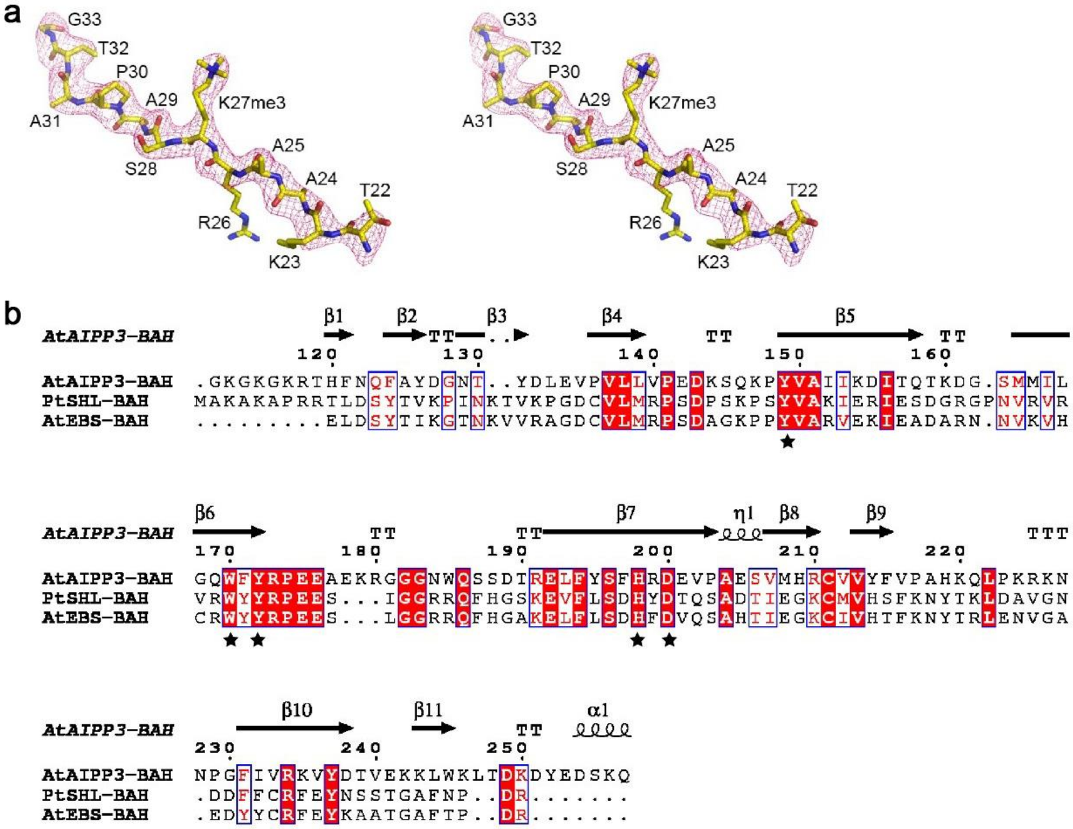


**Supplementary Fig. 7 Structural analysis of the AIPP3 BAH-H3K27me3 complex.** **a,** A stereo view of the SIGMAA-weighted 2Fo-Fc electron density map of the H3K27me3 peptide. **b,** Structure-based sequence alignment of the BAH domains from *Arabidopsis* AIPP3 (AtAIPP3), *Populus trichocarpa* SHL (PtSHL), and *Arabidopsis* EBS (AtEBS) with the secondary structures of AtAIPP3 BAH showing on the top of the alignment. The conserved H3K27me3 peptide-interacting residues are highlighted by stars on the bottom.

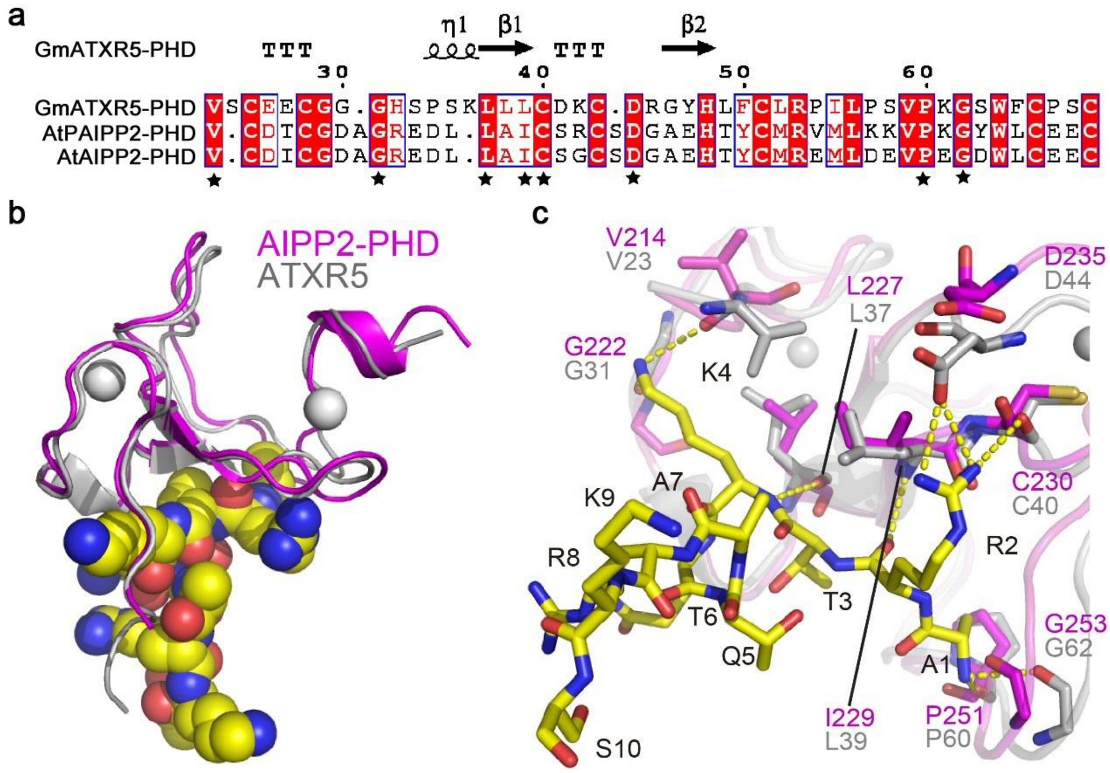


**Supplementary Fig. 8 Structural analysis of the AIPP2 PHD-H3 modeled complex.** **a,** Structure-based sequence alignment of the PHD fingers from *Arabidopsis* AIPP2 and PAIPP2 (AtAIPP2 and AtPAIPP2) and the modeling template *Glycine max* ATXR5 (GmATXR5) with the secondary structures of GmATXR5 showing at the top of the alignment. The conserved H3 peptide-interacting residues are highlighted with stars on the bottom. **b,** The overall modeled structure of the AIPP2 PHD finger (in magenta) in complex with the H3 peptide (in the space-filling model) is overlaid with the structure of the model template ATXR5 PHD finger (in silver). **c,** The detailed interaction between the AIPP2 PHD finger and the H3 peptide with the interacting residues highlighted in the stick model and the hydrogen bonds highlighted in dashed yellow lines. The corresponding residues of the model template ATXR5 PHD finger are overlain, showing that almost all the peptide-binding residues are conserved.

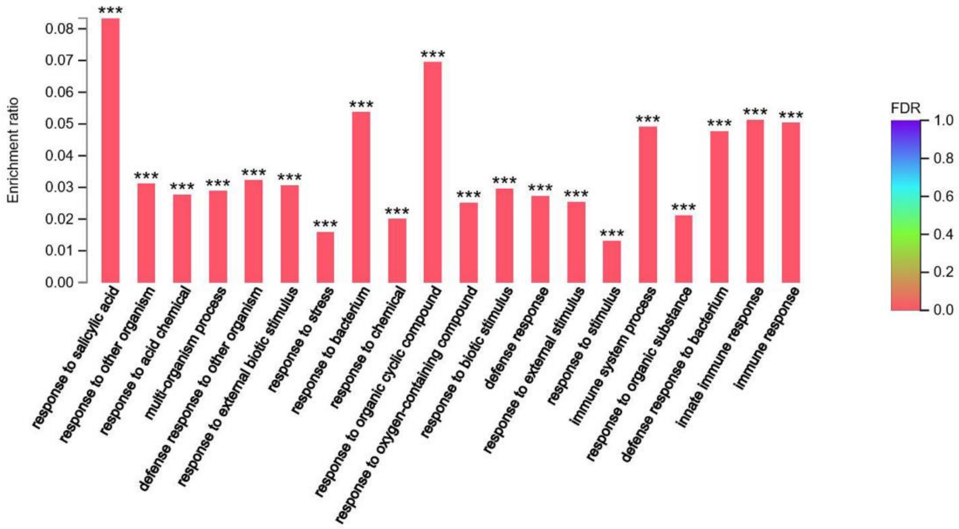


**Supplementary Fig. 9 FDR analysis of commonly up-regulated genes in the mutants of the BAH-PHD-CPL2 complex.** Commonly up-regulated genes were assigned into different GO terms (x-axis). The y-axis indicates the ratio of the up-regulated gene number and the number of genes annotated in this pathway. The color indicates significance of enrichment. ***, p value < 0.001

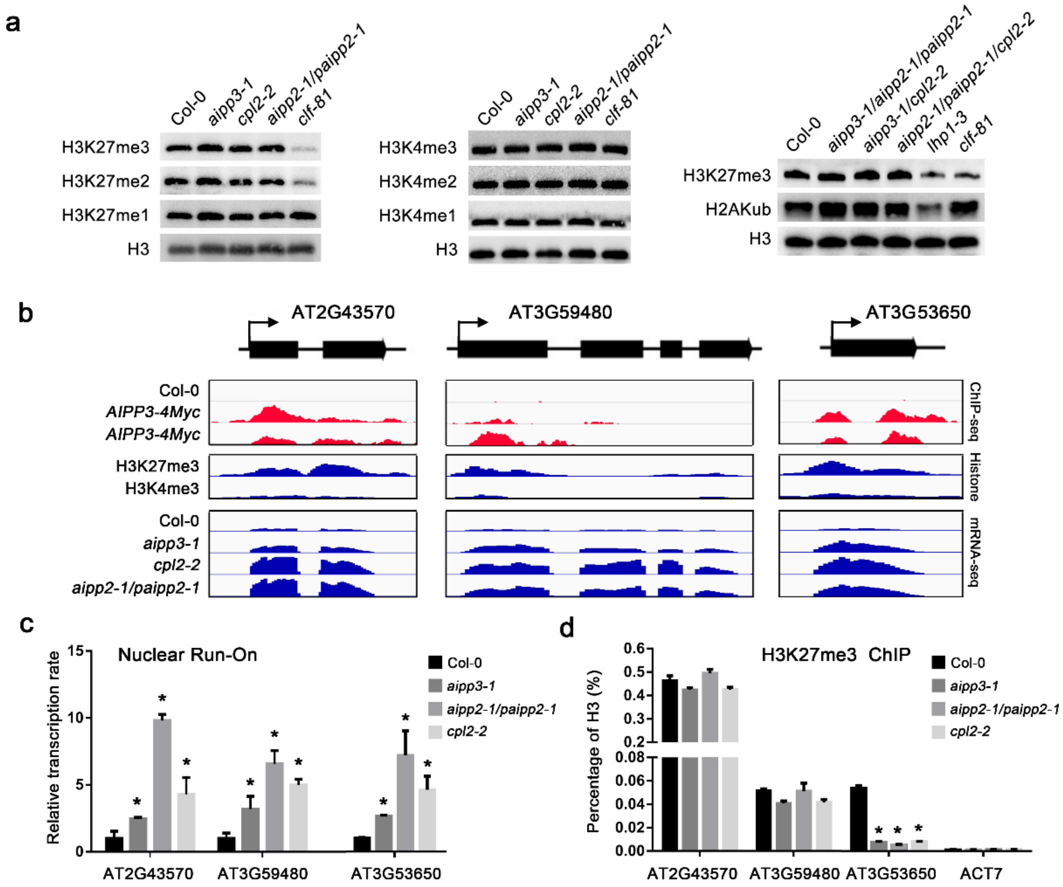


**Supplementary Fig. 10 The effects of BPC dysfunctions on the deposition of different histone marks. a** The immunoblotting results showing the accumulation of the H3K27me1/2/3, H3K4me1/2/3 and H2Kub marks in the selected mutants. The H3 levels were determined to use as the loading controls. **b** Snapshots of IGV of AIPP3 ChIP-seq (upper panel), histone ChIP (middle panel) and mRNA-seq (lower panel) showing the distribution patterns of AIPP3, H3K27me3 and H3K4me3 at selected target genes, and the relative expression of these genes in different genotypes. **c** Nuclear Run-On showing the relative Pol II transcription rate at the selected target genes in Col-0 and *bpc* mutants. The relative transcription rate was normalized to *ACT2*. The Data are the means ±S.D. of three biological repeats. *, p-value<0.01. **d** ChIP-qPCR results showing the relative occupancy of H3K27me3 at selected target genes in Col-0 and *bpc* mutants. The occupancy was first normalized to histone H3. The Data are the means ±S.D. of three biological repeats. *, p-value<0.01.

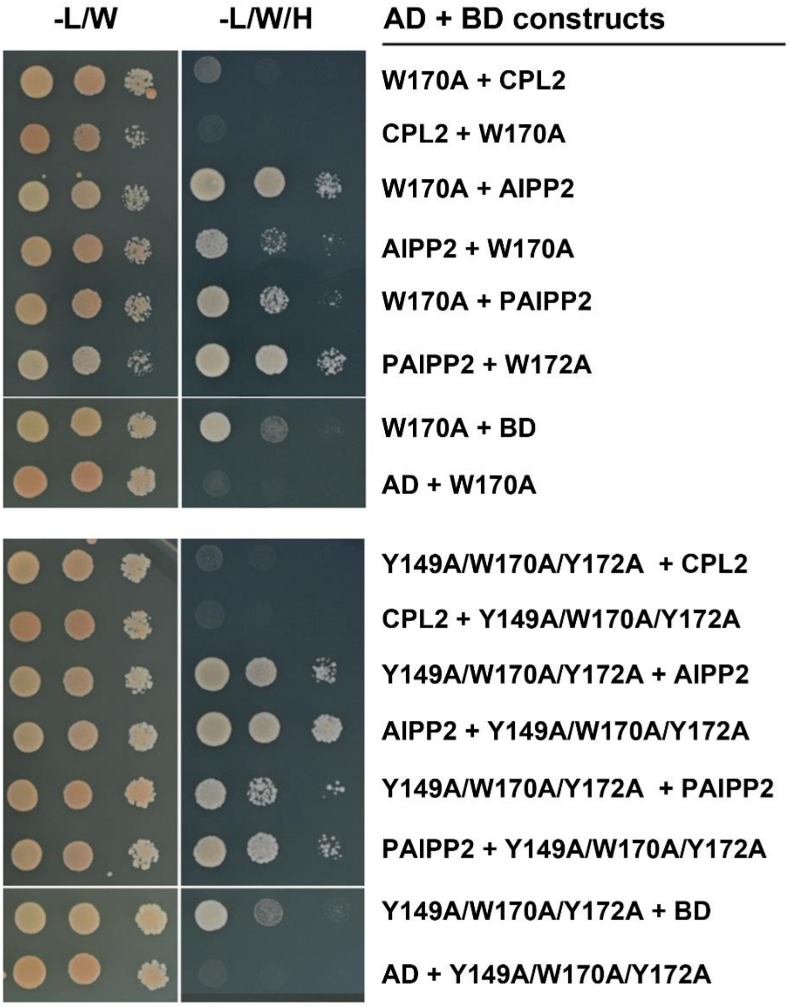


**Supplementary Fig. 11 The 170A and Y149A/W170A/Y172A mutations of AIPP3 did not affect its interaction with AIPP2 and PAIPP2.** Y2H results showing the reciprocal interactions within the tested proteins. Yeast cultures with different protein combinations on SD-LW and SD-LWH are shown.

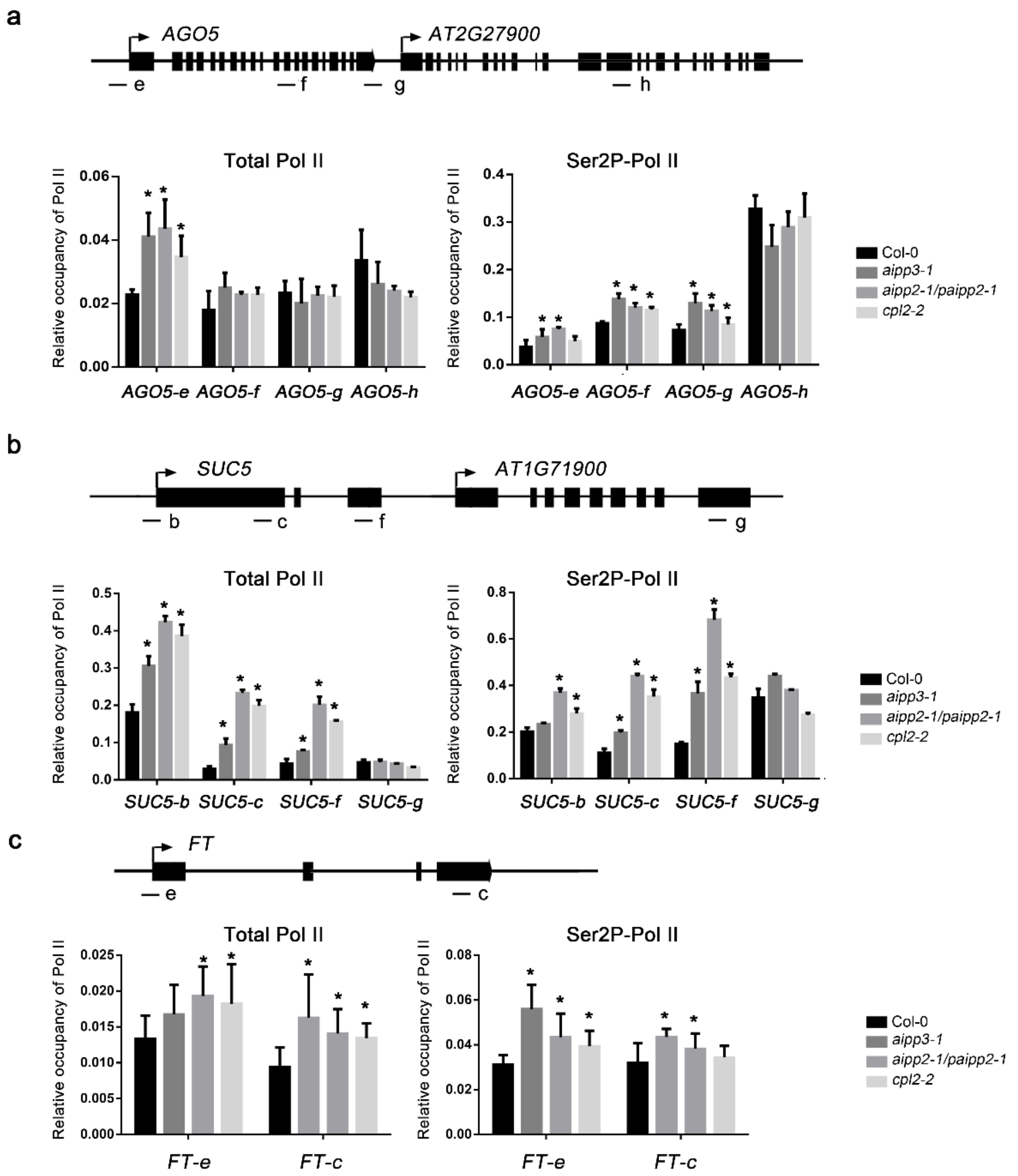


**Supplementary Fig. 12 Relative occupancy of unphosphorylated and Ser2P-Pol II at selected target genes.** ChIP-qPCR showing the relative occupancy of unphosphorylated (total) and Ser2P-Pol II at *AGO5* (a), *SUC5* (b) and *FT* (c) loci. The occupancy was normalized to *ACT7*. The lowercase letters represent positions of ChIP-qPCR primers. The Data are the means ±S.D. of three biological repeats. Significance analysis (t-test) was performed and * represent p-value <0.01.

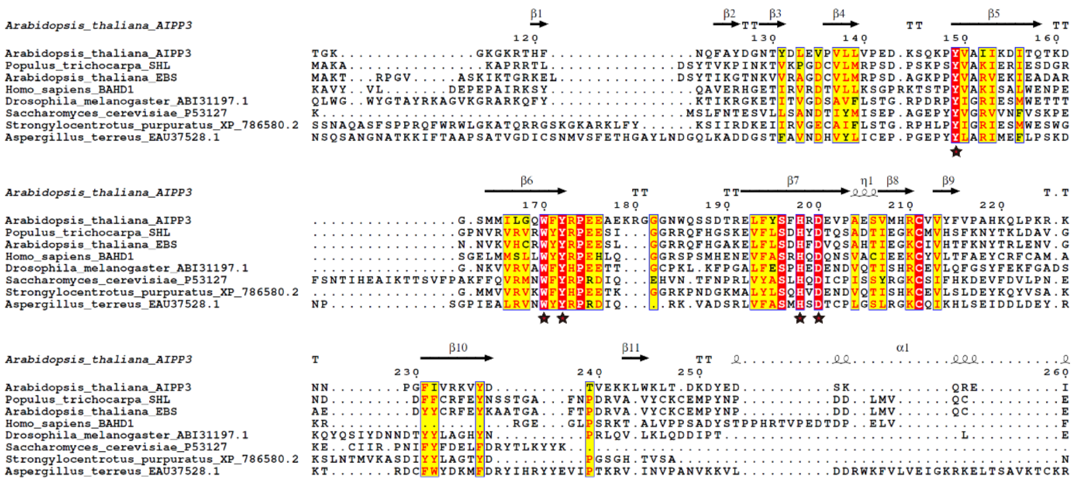


**Supplementary Fig. 13 A structure-based sequence alignment of potential H3K27me3 reading BAH domains from different species.** The secondary structure of Arabidopsis AIPP3 BAH domain is on the top of the alignment. The conserved aromatic cage residues for methyl-lysine binding and histidine and aspartic acid residues for H3P30 binding are marked with stars.

**Supplementary Table 1**

**IP-MS analysis (Supplementary Data 1)**

**Supplementary Table 2**

**Data collection and refinement statistics**

| | AIPP3-BAH-H3K27me3 |
|---|---|
| **Data collection** | |
| Beamline | SSRF-BL19U1 |
| Space group | $P3_121$ |
| Wavelength (Å) | 0.9789 |
| Cell dimensions | |
| $a$=$b$, $c$ (Å) | 78.6, 72.7 |
| Resolution (Å) | 50.0-2.4 (2.44-2.40)[a] |
| $R_{merge}$ | 0.082 (1.121) |
| $I/\sigma I$ | 57.4 (2.8) |
| Completeness (%) | 100.0 (100.0) |
| Redundancy | 19.1 (18.7) |
| **Refinement** | |
| $R_{work}$ / $R_{free}$ | 0.219 / 0.266 |
| No. reflections | 10,440 |
| No. atoms | 1,408 |
| Protein / Peptide | 1,314 / 83 |
| Water / Tris | 3 / 8 |
| $B$-factors (Å$^2$) | 90.7 |
| Protein / Peptide | 90.7 / 92.0 |
| Water / Tris | 63.3 / 93.6 |
| R.m.s. deviations | |
| Bond lengths (Å) | 0.008 |
| Bond angles (°) | 0.955 |

[a] Highest-resolution shell is shown in parentheses.

### Supplementary Table 3 Commonly up-regulated genes in BPC complex mutants

| Gene id | *Col-0* | *Col-0* | *aipp3-1* | *aipp3-1* | *cpl2-2* | *cpl2-2* | *aipp2-1/paipp2-1* | *aipp2-1/paipp2-1* | *aipp2-1* | *aipp2-1* | *paipp2-1* | *paipp2-1* | *clf-81* | *clf-81* | *lhp1-3* | *lhp1-3* |
|---|---|---|---|---|---|---|---|---|---|---|---|---|---|---|---|---|
| AT5G66630 | 1.49 | 1.06 | 4.13 | 5.91 | 3.37 | 4.26 | 4.26 | 4.58 | 1.37 | 1.54 | 3.24 | 2.1 | 0.86 | 0.78 | 0.61 | 0.61 |
| AT5G64810 | 0.4 | 0.58 | 3.08 | 2.04 | 4.26 | 5.32 | 6.41 | 4.96 | 0.55 | 0.6 | 0.9 | 0.82 | 1.6 | 2.07 | 1.42 | 1.61 |
| AT5G64510 | 1.79 | 2.37 | 6.59 | 4.99 | 5.52 | 5.61 | 5.47 | 5.14 | 2.29 | 3 | 3.02 | 2.09 | 2.17 | 1.67 | 1.8 | 1.31 |
| AT5G60900 | 0.5 | 0.95 | 3.58 | 3.4 | 4.7 | 5.24 | 12.9 | 9.54 | 0.85 | 1.3 | 1.12 | 0.59 | 0.85 | 1.09 | 0.3 | 0.45 |
| AT5G59670 | 0.64 | 0.56 | 2.06 | 1.81 | 2.89 | 1.96 | 12.88 | 6.9 | 0.66 | 0.74 | 1.08 | 0.78 | 0.34 | 0.4 | 0.37 | 0.31 |
| AT5G57550 | 0.79 | 0.87 | 1.87 | 3.06 | 2.26 | 2.24 | 9.12 | 8.91 | 1.12 | 1.28 | 1.16 | 1.43 | 6.59 | 6.96 | 0.58 | 0.98 |
| AT5G54610 | 0.8 | 0.92 | 5.57 | 6.16 | 9.88 | 10.03 | 24.2 | 16.75 | 1.45 | 3.09 | 1.56 | 2.17 | 1.68 | 1.69 | 0.61 | 0.46 |
| AT5G54190 | 0.25 | 0.22 | 40.3 | 31.47 | 44.38 | 36.22 | 52.64 | 66.82 | 1.13 | 1.59 | 5.72 | 5.97 | 0.7 | 0.47 | 0.58 | 0.25 |
| AT5G52760 | 0.94 | 2.21 | 15.9 | 7.35 | 9.19 | 9.68 | 60.76 | 20.07 | 1.71 | 4.45 | 3.07 | 1.2 | 2.39 | 3.84 | 0.33 | 0.46 |
| AT5G52750 | 2.73 | 4.3 | 25.55 | 10.77 | 16.2 | 12.7 | 25.6 | 28.22 | 4.21 | 8.11 | 6.53 | 3.62 | 9.13 | 9.38 | 4.3 | 4.25 |
| AT5G52300 | 0.13 | 0.07 | 2.07 | 1.04 | 2.04 | 1.56 | 5.09 | 4.38 | 0.13 | 0.3 | 0.94 | 0.54 | 0.11 | 0.04 | 0.41 | 0.35 |
| AT5G50790 | 0.96 | 0.8 | 3.69 | 3.99 | 6.13 | 4.77 | 5.36 | 3.16 | 1.07 | 1.04 | 2.38 | 1.88 | 19.12 | 21.83 | 11.13 | 12.13 |
| AT5G47600 | 0.28 | 0.97 | 18.66 | 17.45 | 32.87 | 34.42 | 42.73 | 67.65 | 1.18 | 0.89 | 2.86 | 3.34 | 0.68 | 0.86 | 0.88 | 1.06 |
| AT5G45990 | 0.35 | 0.36 | 1.98 | 2.1 | 2.48 | 1.95 | 3.98 | 2.77 | 0.68 | 0.6 | 1.04 | 0.85 | 0.57 | 0.55 | 0.8 | 0.7 |
| AT5G45380 | 3.48 | 5 | 10.36 | 11.93 | 9.3 | 11.97 | 9.26 | 11.23 | 5.06 | 6.31 | 7.66 | 5.91 | 2.75 | 2.34 | 4.76 | 5.01 |
| AT5G44120 | 0.03 | 0.04 | 2.48 | 7.48 | 4.17 | 23.64 | 38.72 | 80.3 | 0.13 | 0.09 | 26.97 | 7.22 | 0.11 | 0.07 | 0.28 | 0.27 |
| AT5G42830 | 2.78 | 3.57 | 8.96 | 10.08 | 8.4 | 11.89 | 30.5 | 23.26 | 3.36 | 5.57 | 5.75 | 3.87 | 7.87 | 7.87 | 7.63 | 7.09 |
| AT5G42530 | 216.24 | 287.47 | 622.75 | 734.49 | 714.59 | 722.95 | 3272.63 | 1215.44 | 295.03 | 323.76 | 419.72 | 409.43 | 162.7 | 209.11 | 83.38 | 80.29 |
| AT5G41750 | 0.15 | 0.05 | 0.75 | 0.31 | 0.17 | 0.25 | 2.58 | 0.67 | 0.1 | 0.23 | 0.08 | 0.07 | 0.24 | 0.23 | 0.09 | 0.09 |
| AT5G38240 | 0.27 | 0.36 | 2.07 | 1.62 | 1.97 | 1.86 | 4.82 | 2.59 | 0.61 | 0.66 | 0.9 | 0.68 | 1.13 | 1.02 | 0.37 | 0.4 |
| AT5G35940 | 2.05 | 3.19 | 14.78 | 13.43 | 8.76 | 8.33 | 15.89 | 9.49 | 3.47 | 3.82 | 5.72 | 4.79 | 21.7 | 28.19 | 3.47 | 4.06 |
| AT5G27420 | 2.19 | 2.24 | 22.52 | 9.06 | 7.62 | 7.33 | 4.52 | 14.01 | 3.49 | 3.99 | 3.51 | 5.55 | 7.57 | 6.92 | 4.1 | 3.81 |
| AT5G25440 | 3.94 | 5.44 | 15.32 | 14.94 | 12.57 | 12.8 | 23.29 | 16.55 | 5.6 | 7.19 | 6.14 | 5.38 | 5.68 | 6.36 | 4.56 | 4.07 |
| AT5G25260 | 0.52 | 0.86 | 2.75 | 3.05 | 4.74 | 7.76 | 6.54 | 3.88 | 0.96 | 1.63 | 1.73 | 0.26 | 2.02 | 1.43 | 0.52 | 0.27 |
| AT5G25250 | 6.88 | 6.6 | 44.29 | 38.85 | 29.26 | 32.21 | 43.86 | 45.32 | 8.61 | 13.88 | 11.9 | 6.42 | 9.16 | 8.74 | 1.74 | 1.61 |
| AT5G24600 | 0.72 | 0.66 | 2.78 | 2.53 | 3.19 | 3.92 | 6.91 | 4.55 | 0.56 | 0.93 | 1.34 | 0.77 | 1.39 | 1.81 | 0.95 | 0.72 |
| AT5G24530 | 15.03 | 14.32 | 53.82 | 48.24 | 64.49 | 73.53 | 87.09 | 54.78 | 20.7 | 27.9 | 17.2 | 15.97 | 18.17 | 17.4 | 13.55 | 13.09 |
| AT5G24210 | 15.21 | 19.97 | 47.4 | 44.36 | 54.59 | 56.87 | 115.58 | 50.2 | 25.62 | 34.94 | 17.3 | 11.88 | 29.25 | 29.43 | 12.92 | 12.38 |
| AT5G20230 | 16.32 | 10.44 | 72.2 | 51.77 | 59.87 | 92.49 | 70.9 | 75.17 | 23.23 | 35.43 | 28.28 | 11.82 | 65.48 | 80.5 | 16.77 | 17.04 |
| AT5G14470 | 0.68 | 0.45 | 5.6 | 6.08 | 6.63 | 8.07 | 8.95 | 6.97 | 1 | 1.38 | 1.51 | 1.73 | 0.74 | 0.23 | 0.66 | 0.84 |
| AT5G11920 | 2.32 | 2.69 | 6.93 | 6.57 | 7.34 | 8.26 | 11.12 | 9.36 | 2.61 | 2.27 | 3.85 | 4.07 | 7.76 | 9.02 | 3.39 | 3.09 |
| AT5G07770 | 3.26 | 3.05 | 7.84 | 8.69 | 7.3 | 8.81 | 11.28 | 7.9 | 3.42 | 4.15 | 4.51 | 4.34 | 1.97 | 1.56 | 2.39 | 2.66 |
| AT5G03350 | 4.06 | 8.33 | 29.52 | 27.21 | 32.84 | 36.08 | 99.55 | 79.9 | 4.31 | 10.94 | 6.89 | 6.22 | 11.74 | 21.95 | 4 | 3.9 |
| AT5G02895 | 0 | 0 | 0 | 0 | 0 | 0 | 0 | 0 | 0 | 0 | 0 | 0 | 0 | 0 | 0 | 0 |
| AT5G02490 | 3.52 | 4.71 | 14.35 | 16.75 | 13.04 | 16.74 | 10.05 | 10.52 | 4.15 | 5.38 | 6.01 | 4.91 | 11.79 | 9.69 | 6.06 | 5.75 |
| AT4G36880 | 2.01 | 2.65 | 9.81 | 11.51 | 6.77 | 8.03 | 16.66 | 10.03 | 2.07 | 2.84 | 6.07 | 5.29 | 2.09 | 2.02 | 5.13 | 4.27 |
| AT4G35180 | 1.37 | 0.84 | 4.29 | 4.12 | 3.21 | 4.83 | 13.72 | 7.61 | 1.7 | 2.26 | 1.12 | 0.28 | 2.13 | 2.48 | 0.68 | 0.5 |
| AT4G33930 | 0.24 | 0.14 | 3.43 | 2.7 | 6 | 9.31 | 34.44 | 19.02 | 0.32 | 0.21 | 1.24 | 0.62 | 0.29 | 0.07 | 0.22 | 0.29 |
| AT4G33120 | 0.47 | 0.5 | 3.21 | 3.28 | 4.77 | 5.26 | 7.84 | 5.38 | 0.78 | 0.55 | 2.1 | 2.3 | 2.18 | 2.86 | 0.53 | 0.88 |
| AT4G32480 | 11.04 | 17.94 | 37.62 | 32.71 | 36.76 | 31.81 | 52.73 | 37.47 | 16.15 | 20.52 | 24.04 | 19.38 | 64.49 | 63.96 | 35 | 35.96 |
| AT4G31520 | 0.12 | 0.39 | 1.54 | 1.5 | 3 | 4.1 | 4.41 | 3.39 | 0.21 | 0.4 | 0.68 | 0.37 | 0.28 | 0.25 | 0.34 | 0.25 |
| AT4G26200 | 1.3 | 1.89 | 5.94 | 3.96 | 4 | 6.54 | 15.37 | 10.3 | 1.88 | 1.89 | 2.98 | 1.78 | 8.21 | 6.81 | 5.4 | 5.04 |
| AT4G25110 | 4.37 | 5.65 | 15.32 | 15.92 | 17.34 | 20.77 | 25.53 | 25.11 | 5.95 | 6.21 | 6.12 | 5.32 | 2.71 | 2.95 | 2.04 | 2.06 |
| AT4G24340 | 2.69 | 3.1 | 15.16 | 13.69 | 11.4 | 17.81 | 18.6 | 15.59 | 3.44 | 2.22 | 5.91 | 7.77 | 19.46 | 22.49 | 3.95 | 4.39 |
| AT4G23700 | 3.15 | 2.6 | 9.52 | 10.53 | 6.35 | 8.33 | 9.86 | 8.75 | 3.98 | 4.61 | 4.55 | 6.23 | 5.84 | 6.2 | 4.41 | 4.28 |
| AT4G23210 | 1.16 | 1.08 | 3.37 | 4.95 | 2.9 | 5.49 | 4.99 | 3.31 | 1.3 | 2.46 | 1.58 | 1.04 | 5.76 | 4.96 | 1.11 | 1.1 |
| AT4G23150 | 0.26 | 0.79 | 3.08 | 2.79 | 3.28 | 3.88 | 13.29 | 6.1 | 0.61 | 1.11 | 0.71 | 0.25 | 0.14 | 0.29 | 0.03 | 0.12 |
| AT4G21380 | 1 | 1.19 | 3.64 | 3.8 | 3.39 | 5.04 | 7.07 | 5.4 | 1.18 | 1.07 | 1.64 | 1.05 | 2.39 | 2.18 | 0.84 | 0.88 |
| AT4G18253 | 4.67 | 5.64 | 14.47 | 12.15 | 12.32 | 14.32 | 13.36 | 27.53 | 4.66 | 6.3 | 3.89 | 3.14 | 3.69 | 3.12 | 3.61 | 4.13 |
| AT4G14365 | 7.07 | 11.65 | 32.38 | 29.52 | 25.05 | 32.96 | 110.86 | 55.27 | 11.87 | 17.43 | 11.44 | 7.12 | 13.87 | 12.99 | 5.17 | 4.23 |
| AT4G11890 | 2.45 | 3.59 | 15.06 | 16.46 | 14.07 | 21.29 | 26.55 | 15.86 | 4.36 | 5.92 | 5.88 | 2.59 | 4.65 | 4.4 | 0.96 | 1.17 |
| AT4G11170 | 0.15 | 0.21 | 1.3 | 1.19 | 0.91 | 1.79 | 6.68 | 2.79 | 0.17 | 0.4 | 0.51 | 0.08 | 0.41 | 0.46 | 0.16 | 0.18 |
| AT4G10500 | 1.93 | 1.85 | 7.07 | 6.97 | 7.82 | 9.51 | 13.36 | 8.44 | 0.89 | 1.89 | 2.2 | 1.16 | 3.23 | 2.27 | 2.69 | 2.03 |
| AT4G04490 | 0.84 | 1.36 | 4.32 | 4.01 | 3.88 | 5.1 | 12.51 | 6.99 | 0.7 | 1.31 | 1.63 | 0.71 | 1.07 | 1.33 | 0.36 | 0.41 |
| AT4G04415 | 1.04 | 1.36 | 3.25 | 3.27 | 4.28 | 4.98 | 10.33 | 10.27 | 1.33 | 1.54 | 4.13 | 1.98 | 1.07 | 1.15 | 1.18 | 1.16 |
| AT4G03450 | 0.73 | 0.68 | 3.51 | 4.55 | 2.17 | 3.02 | 14.3 | 7.86 | 0.54 | 0.85 | 0.89 | 0.52 | 0.06 | 0.16 | 0.07 | 0.11 |
| AT3G61280 | 1.19 | 1.74 | 5.93 | 5.13 | 6.7 | 8.9 | 29.09 | 12.51 | 1.57 | 1.81 | 2.27 | 1.6 | 1.51 | 2.14 | 0.94 | 1.13 |
| AT3G61190 | 0.94 | 0.84 | 7.53 | 2.28 | 3.87 | 2.94 | 6 | 5.41 | 1.1 | 1.57 | 1.53 | 1.83 | 5.11 | 5.23 | 1.74 | 0.93 |
| AT3G60415 | 15.56 | 20.78 | 41.81 | 48.4 | 62.3 | 79.25 | 126.59 | 82.29 | 18.81 | 34.13 | 16.06 | 12.01 | 20.12 | 23.65 | 8.07 | 7.86 |
| AT3G59480 | 1.83 | 1.91 | 7.05 | 8.79 | 10.64 | 16.36 | 6.82 | 14.18 | 3.17 | 2.04 | 2.67 | 3.94 | 0.92 | 1.28 | 1.5 | 0.5 |
| AT3G57460 | 0.93 | 1.13 | 4 | 4.48 | 5.22 | 7.43 | 10.02 | 7.62 | 1.08 | 2.57 | 1.07 | 0.25 | 0.8 | 1.18 | 0.26 | 0.12 |
| AT3G56710 | 1.17 | 2.15 | 11.82 | 7.58 | 8.94 | 7.01 | 24.35 | 11.84 | 2.67 | 2.74 | 3.26 | 3 | 2.02 | 2.27 | 1.27 | 1.6 |
| AT3G56400 | 11.21 | 8.98 | 50.15 | 42.56 | 50.79 | 44.56 | 55.94 | 31.93 | 16.47 | 29.2 | 15 | 12.3 | 10.05 | 9.74 | 3.18 | 2.95 |
| AT3G53650 | 0 | 0 | 0 | 0 | 0 | 0 | 0 | 0 | 0 | 0 | 0 | 0 | 0 | 0 | 0 | 0 |
| AT3G50470 | 0.98 | 1.02 | 3.89 | 4.4 | 5.63 | 4.78 | 13.74 | 8.33 | 1.77 | 1.68 | 1.88 | 1.17 | 1.01 | 1.59 | 0.62 | 0.49 |
| AT3G48850 | 0.41 | 0.8 | 2.53 | 3.72 | 2.45 | 4.15 | 4.24 | 5.34 | 0.44 | 0.76 | 0.91 | 0.82 | 2.39 | 5.14 | 1.4 | 1.42 |
| AT3G48390 | 0.69 | 0.74 | 4.44 | 6.07 | 6.29 | 6.69 | 7.56 | 7.48 | 1.44 | 1.73 | 2.43 | 2.71 | 1.26 | 1.11 | 1.17 | 1.18 |
| AT3G47540 | 4.41 | 4.95 | 11 | 11.23 | 16.17 | 19.77 | 19.62 | 28.91 | 4.81 | 5.66 | 7.29 | 6.84 | 7.77 | 12.85 | 5.38 | 5.11 |
| AT3G46090 | 0.79 | 1.33 | 12.88 | 6.16 | 12.03 | 8.46 | 17.41 | 9.07 | 3.31 | 6.19 | 2 | 2.01 | 10.52 | 11.49 | 5.59 | 3.79 |
| AT3G46080 | 0.59 | 0.98 | 6.74 | 3.9 | 6.33 | 7.16 | 2.85 | 4.14 | 1.25 | 1.63 | 1.65 | 0.4 | 8.21 | 10.16 | 2.53 | 2.87 |
| AT3G45860 | 1.9 | 2.21 | 8.69 | 7.56 | 13.42 | 13.79 | 37.28 | 17.45 | 1.9 | 3.48 | 2.59 | 2.01 | 0.28 | 0.14 | 0.82 | 0.83 |
| AT3G44860 | 1.04 | 1.44 | 4.76 | 5.15 | 2.99 | 6.38 | 6.34 | 23.34 | 0.93 | 1.81 | 1.42 | 0.92 | 35.68 | 40.18 | 1.64 | 1.69 |
| AT3G44400 | 1.48 | 1.75 | 5.25 | 5.19 | 3.49 | 4.09 | 16.37 | 7.55 | 2.37 | 2.86 | 2.84 | 2.49 | 2.22 | 2.73 | 2.59 | 2.42 |
| AT3G29000 | 0.64 | 0.88 | 17.65 | 5.37 | 6.34 | 3.31 | 25.92 | 9.5 | 0.96 | 1.6 | 1.58 | 1.79 | 4.77 | 7.04 | 2.78 | 2.48 |
| AT3G28510 | 0.19 | 0.48 | 2.88 | 4.63 | 2.91 | 4.38 | 13.3 | 2.83 | 0.38 | 0.71 | 0.39 | 0.39 | 1.33 | 1.28 | 2.16 | 1.6 |
| AT3G25010 | 0.08 | 0.11 | 0.2 | 0.21 | 0.08 | 0.03 | 0.05 | 0.1 | 0.11 | 0.1 | 0.09 | 0.03 | 0.01 | 0.04 | 0.17 | 0.1 |
| AT3G22235 | 56.49 | 44.81 | 145.25 | 141.06 | 192.26 | 150.43 | 224.38 | 243.69 | 38.66 | 69.14 | 42.36 | 40.74 | 37.9 | 29.68 | 5.41 | 5.37 |

| | | | | | | | | | | | | | | | | |
|---|---|---|---|---|---|---|---|---|---|---|---|---|---|---|---|---|
| AT3G22231 | 0 | 0 | 0.06 | 0.06 | 0.18 | 0.12 | 0.2 | 0.16 | 0.06 | 0 | 0.06 | 0 | 0.05 | 0 | 0 | 0 |
| AT3G21370 | 0.02 | 0 | 0.63 | 1.35 | 0.73 | 2.03 | 5.09 | 11.21 | 0.02 | 0.06 | 2.94 | 1.01 | 0.22 | 0.17 | 0.37 | 0.09 |
| AT3G13090 | 0.41 | 0.41 | 1.96 | 1.76 | 3.07 | 4.05 | 4.05 | 3.19 | 0.52 | 0.46 | 1.12 | 1.06 | 0.67 | 0.51 | 0.61 | 0.77 |
| AT3G12580 | 20.6 | 23.16 | 75.81 | 88.9 | 62.88 | 70.66 | 68.93 | 63.15 | 20.4 | 24.51 | 53.38 | 48.33 | 23.25 | 20.8 | 25.82 | 27.54 |
| AT3G12220 | 0.52 | 0.49 | 1.98 | 2.99 | 3.1 | 3.16 | 4.52 | 3.11 | 0.28 | 0.75 | 0.52 | 0.57 | 0.53 | 0.68 | 0.22 | 0.35 |
| AT3G11010 | 3.16 | 4.5 | 10.71 | 9.92 | 10.75 | 12.93 | 66.95 | 29.71 | 3.87 | 5.88 | 5.3 | 3.89 | 2.38 | 1.99 | 2.47 | 2.49 |
| AT3G08870 | 0.38 | 0.59 | 1.52 | 2.19 | 1.51 | 2.09 | 3.69 | 2.42 | 0.6 | 0.84 | 0.84 | 0.69 | 1.33 | 1.44 | 0.77 | 0.71 |
| AT3G05995 | 0.08 | 0.23 | 1.47 | 1.42 | 2.68 | 3.33 | 5.52 | 2.41 | 0.34 | 0.46 | 0.4 | 0.2 | 0.21 | 0.18 | 0.31 | 0.15 |
| AT2G46430 | 3.1 | 4.46 | 9.66 | 10.85 | 8.55 | 10.68 | 24.26 | 17.14 | 4.25 | 5.23 | 5.75 | 4.38 | 7.84 | 8.51 | 4.42 | 4.84 |
| AT2G43570 | 7.51 | 5.69 | 21.15 | 16.16 | 51.98 | 67.57 | 35.75 | 51.22 | 8.08 | 10.51 | 14.37 | 9.69 | 15.77 | 21.42 | 10.3 | 9.91 |
| AT2G41470 | 0.06 | 0.17 | 3.24 | 2.95 | 5.24 | 3.69 | 12.2 | 11.2 | 0.13 | 0.08 | 0.36 | 0.15 | 0.11 | 0.36 | 0.15 | 0.06 |
| AT2G41105 | 0 | 0 | 0 | 0 | 0 | 0 | 0 | 0 | 0 | 0 | 0 | 0 | 0 | 0 | 0 | 0 |
| AT2G40750 | 2.51 | 1.84 | 10.93 | 10.8 | 13.95 | 11.32 | 26.94 | 11.4 | 2.61 | 3.91 | 2.43 | 2.38 | 3.01 | 2.73 | 1.31 | 1.2 |
| AT2G39310 | 64.99 | 53.55 | 183.88 | 216.52 | 201.47 | 286.68 | 254.94 | 222.35 | 64.77 | 69.8 | 181.65 | 216.38 | 143.64 | 113.48 | 116.62 | 113.64 |
| AT2G39210 | 8.95 | 12.1 | 24.61 | 25.27 | 20.82 | 26.59 | 45.03 | 30.97 | 12.07 | 16.5 | 13.14 | 10.89 | 14.51 | 16.16 | 7.43 | 6.65 |
| AT2G35980 | 1.83 | 1.04 | 9.8 | 8.91 | 4.74 | 8.78 | 6.47 | 7.55 | 1.6 | 1.72 | 3.79 | 3.13 | 6.59 | 8.76 | 1.72 | 1.83 |
| AT2G31880 | 16.46 | 22.73 | 68.17 | 51.5 | 53 | 61.12 | 77.73 | 94.4 | 20.51 | 26.82 | 26.46 | 19.48 | 31.84 | 29.01 | 20.63 | 19.92 |
| AT2G30750 | 2.19 | 2.52 | 7.57 | 12.32 | 5.23 | 8.97 | 16.33 | 11.02 | 2.81 | 3 | 3.43 | 2.1 | 24.27 | 28.92 | 6.02 | 5.51 |
| AT2G28400 | 2.08 | 3.51 | 10.03 | 10.17 | 8.58 | 9.5 | 8.28 | 11.3 | 4.16 | 5.35 | 6.67 | 5.35 | 9.04 | 12.83 | 3.94 | 3.44 |
| AT2G27660 | 3.03 | 3.76 | 8.52 | 7.93 | 7.03 | 10.08 | 14.32 | 12.66 | 4.95 | 4.33 | 3.67 | 3.52 | 4.94 | 4.44 | 4.17 | 4.46 |
| AT2G20142 | 0 | 0 | 0.02 | 0.02 | 0 | 0 | 0.1 | 0.01 | 0 | 0 | 0 | 0 | 0 | 0 | 0 | 0 |
| AT2G18690 | 9.55 | 8.11 | 34.69 | 32.18 | 17.26 | 23.24 | 46.19 | 19.59 | 8.26 | 12.23 | 11.52 | 5.68 | 10.55 | 13.97 | 3.12 | 3.22 |
| AT2G18660 | 0.99 | 2.29 | 8.52 | 6.77 | 6.82 | 11.93 | 12.76 | 10.03 | 1.51 | 2.59 | 1.12 | 0.42 | 1.43 | 1.9 | 0.48 | 0.34 |
| AT2G17040 | 1.7 | 2.87 | 12.85 | 6.51 | 12.62 | 11.14 | 23.31 | 14.14 | 2.67 | 5.23 | 4.08 | 1.8 | 4.97 | 4.67 | 2.9 | 2.51 |
| AT2G14610 | 14.36 | 28.04 | 98.54 | 81.68 | 341.67 | 330.33 | 474.11 | 728.14 | 59.33 | 26.72 | 15.28 | 2.29 | 83.69 | 151.59 | 31.65 | 32.99 |
| AT2G14560 | 4.08 | 8.66 | 33.22 | 33.42 | 31.49 | 35.41 | 56.61 | 64.74 | 6.03 | 12.33 | 4.88 | 3.65 | 12.99 | 16.7 | 4.4 | 4.08 |
| AT2G13810 | 0.39 | 0.36 | 2.26 | 2.06 | 4.55 | 5.52 | 4.05 | 3.55 | 0.27 | 0.94 | 0.62 | 0.31 | 0.82 | 0.94 | 0.16 | 0.08 |
| AT2G02930 | 1.12 | 0.66 | 3.86 | 4.09 | 3.72 | 4.57 | 4.73 | 4.45 | 1.56 | 1.72 | 2.34 | 1.37 | 0.51 | 0.16 | 0.67 | 0.36 |
| AT1G80840 | 2.66 | 3.29 | 37.88 | 14.05 | 12.15 | 10.84 | 57.96 | 25.32 | 4.73 | 8.04 | 6.62 | 4.75 | 11.72 | 15.39 | 3.88 | 3.2 |
| AT1G76960 | 2.99 | 6.81 | 17.28 | 16.88 | 16.25 | 20.05 | 24.26 | 31.11 | 6.05 | 10.52 | 5.86 | 4.34 | 5.36 | 5.18 | 2.22 | 1.14 |
| AT1G74870 | 0.34 | 0.43 | 2.47 | 2.29 | 12.34 | 12.82 | 19.74 | 8.95 | 0.92 | 0.72 | 0.93 | 0.73 | 0.42 | 0.28 | 0.35 | 0.47 |
| AT1G73805 | 1.78 | 2.01 | 10.86 | 7.68 | 11.07 | 12.73 | 52.96 | 18.35 | 1.89 | 2.99 | 2.51 | 1.72 | 3.9 | 3.59 | 0.66 | 0.58 |
| AT1G72930 | 2.72 | 4.78 | 8.53 | 7.85 | 9.15 | 8.2 | 17.05 | 20.95 | 4.46 | 5.34 | 5.73 | 4.89 | 11.47 | 11.35 | 9.25 | 8.5 |
| AT1G72910 | 1.7 | 1.23 | 14.42 | 5.88 | 8.98 | 5.13 | 17.98 | 11.31 | 2.86 | 3.05 | 5.1 | 5.03 | 6.37 | 5.79 | 2.82 | 2.25 |
| AT1G72060 | 0 | 0.06 | 0.31 | 0.35 | 0.25 | 0.18 | 0 | 0.29 | 0.13 | 0.05 | 0.06 | 0.18 | 0.23 | 0.62 | 0.2 | 0.23 |
| AT1G71890 | 0.19 | 0.46 | 3.73 | 2.46 | 11.17 | 10.19 | 12.83 | 24.91 | 0.77 | 0.92 | 1.71 | 1.68 | 1.29 | 1.08 | 1.32 | 1.35 |
| AT1G67980 | 0.63 | 0.18 | 6.38 | 4.56 | 10.42 | 12.81 | 6.8 | 9.98 | 1.09 | 1.04 | 2.71 | 1.6 | 0.78 | 0.63 | 0.49 | 0.22 |
| AT1G67370 | 0.52 | 0.76 | 3.97 | 4.39 | 12.83 | 15.33 | 22.85 | 12.64 | 0.85 | 0.99 | 1.46 | 1.53 | 0.44 | 0.56 | 1.17 | 0.74 |
| AT1G63530 | 2.18 | 3.48 | 7.71 | 6.83 | 7.19 | 6.65 | 8.07 | 7.61 | 2.99 | 3 | 3.54 | 3.9 | 3.63 | 3.03 | 3.64 | 3.36 |
| AT1G62290 | 2.32 | 3.86 | 8.96 | 9.73 | 10.51 | 11.46 | 11.57 | 15.33 | 2.79 | 3.25 | 6 | 6.77 | 3.94 | 3.39 | 3.16 | 3.01 |
| AT1G57630 | 0.07 | 0 | 0 | 0 | 0 | 0 | 0 | 0 | 0 | 0 | 0 | 0 | 0 | 0 | 0 | 0 |
| AT1G52690 | 0.57 | 0.81 | 4.22 | 2.68 | 7.95 | 4.35 | 16.9 | 9.5 | 0.41 | 0.84 | 1.92 | 1.79 | 1.24 | 0.91 | 1.35 | 1.34 |
| AT1G52100 | 2.8 | 2.5 | 13.17 | 12.65 | 28.76 | 31.37 | 38.46 | 46.33 | 5.44 | 4.59 | 23.11 | 25.09 | 17.46 | 15.58 | 19.84 | 20 |
| AT1G51270 | 4.46 | 4.43 | 10.73 | 13.15 | 10.18 | 10.16 | 15.62 | 15.39 | 4.61 | 4.74 | 6.5 | 5.61 | 12.08 | 12.31 | 8.72 | 8.63 |
| AT1G35710 | 5.37 | 8.28 | 29.85 | 27.89 | 27.55 | 33.19 | 140.26 | 61.08 | 7.96 | 15.57 | 10.39 | 5.99 | 4.21 | 4.75 | 4.64 | 4.41 |
| AT1G35230 | 1.78 | 2.67 | 6.91 | 7.85 | 15.46 | 19.63 | 11.52 | 13.92 | 2.77 | 5.06 | 4.69 | 1.77 | 10.18 | 9.8 | 1.75 | 1.43 |
| AT1G35210 | 0.98 | 0.93 | 14.03 | 5.1 | 5.49 | 3.14 | 6.34 | 7.81 | 1.58 | 2.96 | 2.93 | 3.69 | 1.02 | 1.2 | 0.88 | 0.42 |
| AT1G33960 | 1.93 | 4.38 | 11.76 | 13.06 | 25.42 | 31.68 | 36.44 | 14.44 | 4.12 | 15.38 | 7.65 | 2.92 | 12.13 | 14.51 | 0.92 | 0.64 |
| AT1G33720 | 1.18 | 1.28 | 3.12 | 3.48 | 4.24 | 4.68 | 13.15 | 5.05 | 1.3 | 2.08 | 1.63 | 1.53 | 3.91 | 3.56 | 4.51 | 3.88 |
| AT1G32960 | 0.52 | 0.55 | 4.9 | 4.18 | 4.33 | 7.04 | 11.43 | 9.13 | 0.43 | 0.47 | 0.84 | 0.61 | 1.81 | 1.45 | 0.18 | 0.24 |
| AT1G30370 | 1.26 | 1.28 | 13.49 | 4.96 | 3.48 | 3.84 | 8.38 | 5.52 | 1.86 | 3.04 | 2.53 | 2.32 | 1.35 | 1.09 | 0.64 | 0.59 |
| AT1G28480 | 0.57 | 0.15 | 5.89 | 4.04 | 3.29 | 5.81 | 5 | 8.43 | 0.79 | 1.31 | 0.87 | 0.93 | 6.08 | 7.3 | 1.75 | 2.08 |
| AT1G27730 | 3.27 | 5.2 | 80.65 | 27.96 | 27.78 | 14.74 | 22.93 | 26.53 | 14.01 | 14.5 | 15.78 | 19.7 | 16.29 | 14.37 | 5.26 | 6.35 |
| AT1G26420 | 0.62 | 1.07 | 4.42 | 4.02 | 3.51 | 6.09 | 9.72 | 7.47 | 1.18 | 1.61 | 1.99 | 0.86 | 1.71 | 2.35 | 1.05 | 0.78 |
| AT1G26390 | 0.79 | 0.44 | 4.22 | 1.81 | 2 | 4.27 | 3.22 | 2.63 | 0.71 | 0.43 | 0.58 | 0.53 | 6.53 | 6.08 | 0.83 | 0.83 |
| AT1G26380 | 0.87 | 0.55 | 7.79 | 5.61 | 3.58 | 4.93 | 26.55 | 17.53 | 1.3 | 1.88 | 3.04 | 1.89 | 2.98 | 5.93 | 0.91 | 0.8 |
| AT1G24147 | 1.45 | 1.09 | 11.16 | 9.26 | 9.59 | 8.09 | 42.58 | 28.97 | 2.3 | 3.6 | 2.53 | 2.4 | 6.17 | 6.92 | 1.5 | 1.3 |
| AT1G24145 | 0.68 | 1.94 | 7.13 | 4.63 | 5.42 | 5.53 | 15.5 | 11.71 | 1.27 | 3.2 | 2.09 | 1.31 | 4.92 | 4.53 | 1.37 | 1.51 |
| AT1G23840 | 0.45 | 0.72 | 2.81 | 2.5 | 2.4 | 3.66 | 9.05 | 5 | 0.79 | 1.42 | 0.86 | 0.69 | 0.67 | 0.73 | 0.47 | 0.67 |
| AT1G21240 | 0.82 | 1.42 | 6.02 | 3.93 | 4.04 | 6.67 | 11.04 | 8.2 | 0.91 | 1.65 | 1.33 | 0.43 | 0.35 | 0.58 | 0.03 | 0.08 |
| AT1G21120 | 0.26 | 0.25 | 1.91 | 1.13 | 0.84 | 1.1 | 1.32 | 3.14 | 0.26 | 0.19 | 0.58 | 0.37 | 0.88 | 1.19 | 1.49 | 1.03 |
| AT1G21100 | 2.4 | 2.05 | 10.18 | 9.28 | 8.61 | 11.45 | 14.56 | 11.02 | 3.95 | 3.4 | 4.88 | 6.15 | 9.12 | 6.6 | 3.3 | 3.5 |
| AT1G19960 | 3.55 | 4.53 | 9.26 | 11.78 | 11.09 | 12.27 | 15.79 | 17.09 | 3.75 | 4.87 | 5.13 | 4.63 | 4.67 | 5.36 | 3.32 | 2.72 |
| AT1G19250 | 0.37 | 0.83 | 4.97 | 6.3 | 6.23 | 11.45 | 5.12 | 7.75 | 0.54 | 1.3 | 2.68 | 0.61 | 0.39 | 0.75 | 0.31 | 0.59 |
| AT1G18830 | 0.03 | 0.06 | 2.4 | 3.25 | 16.13 | 15.71 | 33.21 | 22.93 | 0.14 | 0.21 | 0.74 | 0.83 | 0.1 | 0.08 | 0.18 | 0.31 |
| AT1G16110 | 1.5 | 1.53 | 4.82 | 5 | 4.66 | 4.85 | 11.31 | 6.9 | 1.59 | 2.59 | 2.85 | 2.04 | 2.89 | 3.07 | 1.98 | 1.97 |
| AT1G15520 | 1.76 | 1.23 | 4.93 | 3.68 | 5.06 | 7.38 | 5.32 | 7.48 | 2.02 | 2.77 | 1.64 | 0.9 | 3.38 | 7.47 | 1.15 | 0.97 |
| AT1G15460 | 1.08 | 0.78 | 4.29 | 3.76 | 2.32 | 3.93 | 2.42 | 3.46 | 1.3 | 1.22 | 1.6 | 2.32 | 0.81 | 0.67 | 1.08 | 1.06 |
| AT1G15010 | 4.52 | 3.13 | 9.6 | 11.86 | 12.81 | 17.1 | 15.32 | 12.81 | 3.13 | 5.57 | 3.79 | 3.64 | 7.09 | 6.84 | 2 | 0.97 |
| AT1G14880 | 1.79 | 1.77 | 33.96 | 38.65 | 41 | 53.86 | 42.57 | 39.06 | 3.19 | 7.04 | 8.41 | 0.81 | 1.46 | 0.48 | 0.08 | 0.45 |
| AT1G10340 | 1.35 | 2.52 | 9.12 | 7.89 | 8.63 | 10.59 | 12.64 | 10.19 | 2.33 | 3.22 | 2.21 | 2.11 | 1.8 | 1.47 | 0.81 | 0.53 |
| AT1G09932 | 4.82 | 4.06 | 10.31 | 11.64 | 8.76 | 12.57 | 24.04 | 9.63 | 5.18 | 5.85 | 2.87 | 2.66 | 3.46 | 3.66 | 2.84 | 2.71 |
| AT1G05675 | 0 | 0 | 0 | 0 | 0 | 0 | 0 | 0 | 0 | 0 | 0 | 0 | 0 | 0 | 0 | 0 |
| AT1G02930 | 56.16 | 80.69 | 182.25 | 159.12 | 150.8 | 190.6 | 164.18 | 169.2 | 106.71 | 131.3 | 73.57 | 38 | 169.19 | 210.37 | 38.23 | 36.13 |
| AT1G02920 | 7.83 | 9.84 | 25.61 | 20.46 | 22.18 | 25.32 | 25.76 | 31.92 | 10.69 | 13 | 11.65 | 7 | 18.83 | 18.91 | 5.56 | 5.64 |
| AT1G01680 | 0.14 | 0.53 | 2.23 | 2.51 | 4.59 | 3.75 | 7.61 | 2.84 | 0.14 | 1.1 | 0.95 | 0.45 | 1.52 | 1.01 | 0.31 | 0.06 |
| AT1G01560 | 0.4 | 0.83 | 4.15 | 2.97 | 2.67 | 3.99 | 14.42 | 11.26 | 0.51 | 1.32 | 1.01 | 0.39 | 3.7 | 3.96 | 0.6 | 0.63 |

Note: the numbers indicate the FPKM values in each genotype.

#### Supplementary Table 4. Primers used in this study

| Primer | Sequence (5'-3') | Purpose |
|---|---|---|
| FLC-qF | GCAACGGTCTCATCGAGAAAGCT | RT-qPCR |
| FLC-qR | GATCATCAGCATGCTGTTTCCCAT | |
| FT-qF | GACCTCAGGAACTTCTATACTTTGGTTATG | |
| FT-qR | CTGTTTGCCTGCCAAGCTG | |
| SOC1-qF | GAGAAAGCTCTAGCTGCAGAA | |
| SOC1-qR | CTTGGGCTACTCTCTTCATCAC | |
| AGO5-qF | GAGTAAGCGAAGGGCAGTTTAG | |
| AGO5-qR | CACGAAAGTAACACGAGGAACA | |
| SUC5-qF | ATTGGATGGGTCGTGAAGTG | |
| SUC5-qR | CCTGAACTCCTTGGTCGTAAAG | |
| ACT2-qF | AGGTCCAGGAATCGTTCACA | RT /NRO-qPCR |
| ACT2-qR | GAGTTTGTCACACACAAGTGC | |
| 18SrRNA-NRO-qF | TCCTAGTAAGCGCGAGTCATCA | NRO-qPCR |
| 18SrRNA-NRO-qR | CGAACACTTCACCGGATCAT | |
| AGO5-NRO-qF | GAGTAAGCGAAGGGCAGTTTAG | |
| AGO5-NRO-qR | CACGAAAGTAACACGAGGAACA | |
| AT2G27900-NRO-qF | GAGAAGGAGACATGGACGAAAT | |
| AT2G27900-NRO-qR | GAGTGAGGGAATCTGGAAGAAC | |
| SUC5-NRO-qF | ATTGGATGGGTCGTGAAGTG | |
| SUC5-NRO-qR | CCTGAACTCCTTGGTCGTAAAG | |
| AT1G71900-NRO-qF | GTTGCGGATAAGCAAGCATATAA | |
| AT1G71900-NRO-qR | GTTATCGCAGTGGACCAGAA | |
| FT-NRO-qF | GGCCAAAGAGAGGTGACTAATG | |
| FT-NRO-qR | GGTCTTCTCCACCAATCTCAAC | |
| AT2G43570-F | GTGTCACCGTTCCTTCTAGTG | NRO/ChIP-qPCR |
| AT2G43570-R | GTTGCAGCAAATATGGCTACTG | |
| AT3G53650-F | AGCATCAATGGCTCCGAAA | |
| AT3G53650-R | CTTCTTCTCCGCCTTTGGTAA | |
| AT3G59480-F | CTGCATCTAACGGCGAGAAA | |
| AT3G59480-R | TTGATGAAGCCAGGAGCATC | |
| AGO5a-qF | CCTCCAATCGAATCCACAAGAT | ChIP-qPCR |
| AGO5a-qR | GTCCATAACATAACGTGCCAAATAG | |
| AGO5b-qF | GAAACGTCGGAAGAGGTGAA | |
| AGO5b-qR | AGACGAAGACGCAACAGAAA | |
| AGO5c-qF | GAGTAAGCGAAGGGCAGTTTAG | |
| AGO5c-qR | CACGAAAGTAACACGAGGAACA | |
| AGO5d-qF | GGCACTGGTGTCATGGATTTA | |
| AGO5d-qR | CTCAGGACGATCGGAGATAGAA | |
| AGO5e-qF | CCGGGAATGCAAAGAGATGA | |
| AGO5e-qR | AAACTGGTTACTTGTGTGTATTGTG | |

| | |
|---|---|
| AGO5f-qF | CCTGCTATTCCGTTCATCTCTT |
| AGO5f-qR | GATCCAGTCACATCAGGCAATA |
| AGO5g-qF | TTGAAGGATCTGGCCAACTT |
| AGO5g-qR | TGAGCCAAATCATCGTCCAA |
| AGO5h-qF | GAGAAGGAGACATGGACGAAAT |
| AGO5h-qR | GAGTGAGGGAATCTGGAAGAAC |
| SUC5a-qF | GACCGTCTATTTCACCCTTCTT |
| SUC5a-qR | GTGTGATCTTCTGGGTCTTTCT |
| SUC5b-qF | CTCCAATGTCTCATCTCCTACATC |
| SUC5b-qR | CTCTAACGCCGTAGCATTGT |
| SUC5c-qF | ATTGGATGGGTCGTGAAGTG |
| SUC5c-qR | CCTGAACTCCTTGGTCGTAAAG |
| SUC5d-qF | CCTAGTCTCACAGCAGGAATTA |
| SUC5d-qR | GACCCAGACGAACCCTAAAT |
| SUC5e-qF | GGCAACTTTGGCAAGTATGTT |
| SUC5e-qR | TGTTACGCCAGCCTTGTT |
| SUC5f-qF | CCAATGGGCCTTTCACTCTT |
| SUC5f-qR | AGAACTGGCCTCATATTCCTTAATC |
| SUC5g-qF | GTTGCGGATAAGCAAGCATATAA |
| SUC5g-qR | GTTATCGCAGTGGACCAGAA |
| FTa-qF | GTTCGGACATTGGTAGGTATGG |
| FTa-qR | AAGGGATCCTTCAGGTTAGATAGA |
| FTb-qF | GGCCAAAGAGAGGTGACTAATG |
| FTb-qR | GGTCTTCTCCACCAATCTCAAC |
| FTc-qF | GGAATTCATCGTGTCGTGTTT |
| FTc-qR | GGAAGGCCGAGATTGTAGAT |
| FTd-qF | TGGGATAAATACGAGGAACAACT |
| FTd-qR | GATCTAACCATAACTTGACAGCATAAC |
| FTe-qF | TTCACCGACCCGAGTTAATG |
| FTe-qR | GGTGGTTTCTCTGTGTTGATTG |
| ACT7-qF | CGTTTCGCTTTCCTTAGTGTTAGCT |
| ACT7-qR | AGCGAACGGATCTAGAGACTCACCTTG |
| AtSN1-qF | CCAGAAATTCATCTTCTTTGGAAAAG |
| AtSN1-qR | GCCCAGTGGTAAATCTCTCAGATAGA |
